# *In vivo* translatome profiling reveals early defects in ribosome biology underlying SMA pathogenesis

**DOI:** 10.1101/103481

**Authors:** Paola Bernabò, Toma Tebaldi, Ewout JN Groen, Fiona M Lane, Elena Perenthaler, Francesca Mattedi, Helen J Newbery, Haiyan Zhou, Paola Zuccotti, Valentina Potrich, Francesco Muntoni, Alessandro Quattrone, Thomas H Gillingwater, Gabriella Viero

**Author notes:** These authors contributed equally to this work (Paola Bernabò, Toma Tebaldi, Ewout JN Groen, Fiona M Lane). = corresponding-authors. Correspondence and request for materials should be addressed to G.V. and T.H.G. **Abbreviations** ASO: antisense oligonucleotide FRP: fraction of ribosomes engaged on polysomes NGS: next generation sequencing SMA: spinal muscular atrophy SMN: survival motor neuron protein snRNP: small nuclear ribonucleoprotein TE: translation efficiency.

## Abstract

**Background:** Genetic alterations impacting on ubiquitously expressed proteins involved in mRNA metabolism often result in neurodegenerative conditions, with increasing evidence suggesting that translational defects can contribute to disease. Spinal Muscular Atrophy (SMA) is a neuromuscular disease caused by low levels of SMN protein, whose role in disease pathogenesis remains unclear.

**Results:** By determining in parallel the *in vivo* transcriptome and translatome in SMA mice we identified a robust decrease in translational efficiency, arising during early stages of disease. Translational defects affected translation-related transcripts, were cell autonomous, and were fully rescued after treatment with antisense oligonucleotides to restore SMN levels. Defects in translation were accompanied by a decrease in the number of ribosomes in motor neurons *in vivo.*

**Conclusion:** Our findings suggest that neuronal tissues and cells are particularly sensitive to perturbations in translation during SMA, and identify ribosome biology as an important, yet largely neglected, factor in motor neuron degeneration.

## Background

Neurons are highly specialised cells, reliant on precise spatial and temporal control of translation to maintain their unique anatomy and physiology. Local control of protein synthesis allows neurons to regulate axonal outgrowth and growth cone dynamics during development, and to control the complex cellular processes required for maintaining neuronal homeostasis throughout the life-span [1]. It is therefore not surprising that dysregulation of translation has been linked to the pathogenesis of several neurological diseases [2–7]. Genetic studies of familial and sporadic forms of motor neuron diseases have identified mutations in genes encoding RNA binding proteins (RBPs) or in their interactors as active contributors to disease pathogenesis [8–11]. It remains unclear, however, why defects in ubiquitously expressed components of RNA metabolism lead to defects that are largely restricted to selected cell populations, and the extent to which changes in translation play a causative role in neurodegeneration *in vivo* remains to be fully determined.

Spinal muscular atrophy (SMA) is the most common genetic cause of infant mortality, with an incidence of around 1 in 6,000-10,000 live births [12, 13]. SMA is primarily associated with a loss of lower motor neurons from the spinal cord, leading to muscle atrophy and, eventually, paralysis. In the most common and severe cases of SMA (type I), onset typically occurs around 6 months of age and mortality before two years [13]. SMA is caused by low levels of full-length survival of motor neuron protein (SMN). Humans have two copies of the gene encoding SMN; *SMN1* and *SMN2*. Approximately 95% of SMA cases are caused by homozygous deletion of *SMN1*, with a smaller number caused by discrete mutations within the gene. *SMN1* and *SMN2* are 99.9% identical, with just one base pair difference [14], resulting in skipping of exon 7 in around 85-90% of *SMN2* transcripts. Without exon 7 the *SMN2*-derived protein is quickly degraded and, therefore, only low levels of functional SMN protein remain [15, 16].

Several lines of evidence suggest that SMN is involved in the development and maintenance of motor neuron growth cones, axon, and neuromuscular junctions [17–21] mediated, at least in part, through its role in regulating ubiquitin homeostasis [22, 23]. In concert with other RNA binding proteins, SMN forms complexes both in the nucleus and the cytoplasm of neurons [24, 25]. As such, SMN protein has housekeeping roles in the assembly of ribonucleoprotein (RNP) complexes, including the U3 snoRNA [26] required for 18S ribosomal RNA biogenesis [27] and the small nuclear RNP (snRNP) required for processing of histone pre-mRNAs [28]. In addition, SMN localises to dendrites, synapses and axons *in vivo* and *in vitro* [23, 29, 30] where it is part of messenger RNP (mRNP) complexes with well-known roles in the transport of mRNA [31, 32]. These cellular processes are tightly linked to local translation, suggesting that SMN may play a pivotal role in its regulation. Indeed, SMN has previously been shown to associate with polyribosomes both *in vitro* [33] and in the rat spinal cord [29]. Moreover, components of the translational machinery are mislocalized in SMN-depleted cells [34] and the localisation and translation of specific mRNAs is altered in primary neurons derived from mouse models of SMA [17, 35, 36]. However, a direct link between SMN and the regulation of translation *in vivo* has not yet been demonstrated.

In the current study, we used polysomal profiling to directly investigate the role of SMN in protein translation, *in vitro* and *in vivo*, in an established mouse model of severe SMA at both early and late-symptomatic stages. We show that SMN depletion leads to widespread perturbations in translation, including defects in the fraction of ribosomes on polysomes and in the recruitment of mRNAs to polysomes. These defects are cell-autonomous, directly associated with SMA disease progression, and dependent upon SMN protein levels. Sequencing of mRNAs associated to polysomes led to the identification of specific RNAs differentially translated in SMA. Functional annotation analysis of mRNAs with reduced translation efficiency revealed an enrichment for ribosomal components and proteins involved in translation; consistently, we observed a reduced number of axonal ribosomes in SMA motor neurons. Our findings, therefore, identify a key role for SMN in regulating translation and ribosome biology *in vivo*.

## Results

### Polysomal profiling reveals translational defects in in SMA

To test the hypothesis that low levels of SMN lead to defects in translation, we initially performed polysomal profiling on a set of tissues (brain, spinal cord, kidney and liver) harvested from the severe ‘Taiwanese’ SMA mouse model [23, 37] at pre-, early- and late-symptomatic stages of disease. We did not include disease end-stage mice in these initial experiments, as the severity of the phenotype at this time-point can likely lead to secondary, non-specific changes. Polysomal profiling is a powerful way of gathering a large amount of information about the translational state of biological systems, both *in vitro* and *in vivo* [38–41]. Under both physiological and diseased conditions, it is possible to determine, : 1) the fraction of ribosomes engaged on polysomes, revealing the overall level and efficiency of translation [38]; 2) the protein co-sedimentation profile with ribosomes and polysomes [2, 42], and 3) the transcriptome-wide analysis of RNAs associated with polysomes (i.e. the translatome, POL-Seq) by next generation sequencing (NGS) [43] (Fig. 1a). Graphical representations of typical polysome profiles generated by this approach are shown in Fig. 1b for brain and in Fig. 1c for spinal cord. The first peak contains free cytosolic light components (RNPs) and the subsequent peaks include ribosomal subunits (40S and 60S) and monosomes (80S), all associated with non-translating particles. The remaining peaks of the profile represent polysomes, which sediment with high sucrose concentrations and contain the RNAs associated with ribosomes.

**Figure 1.**
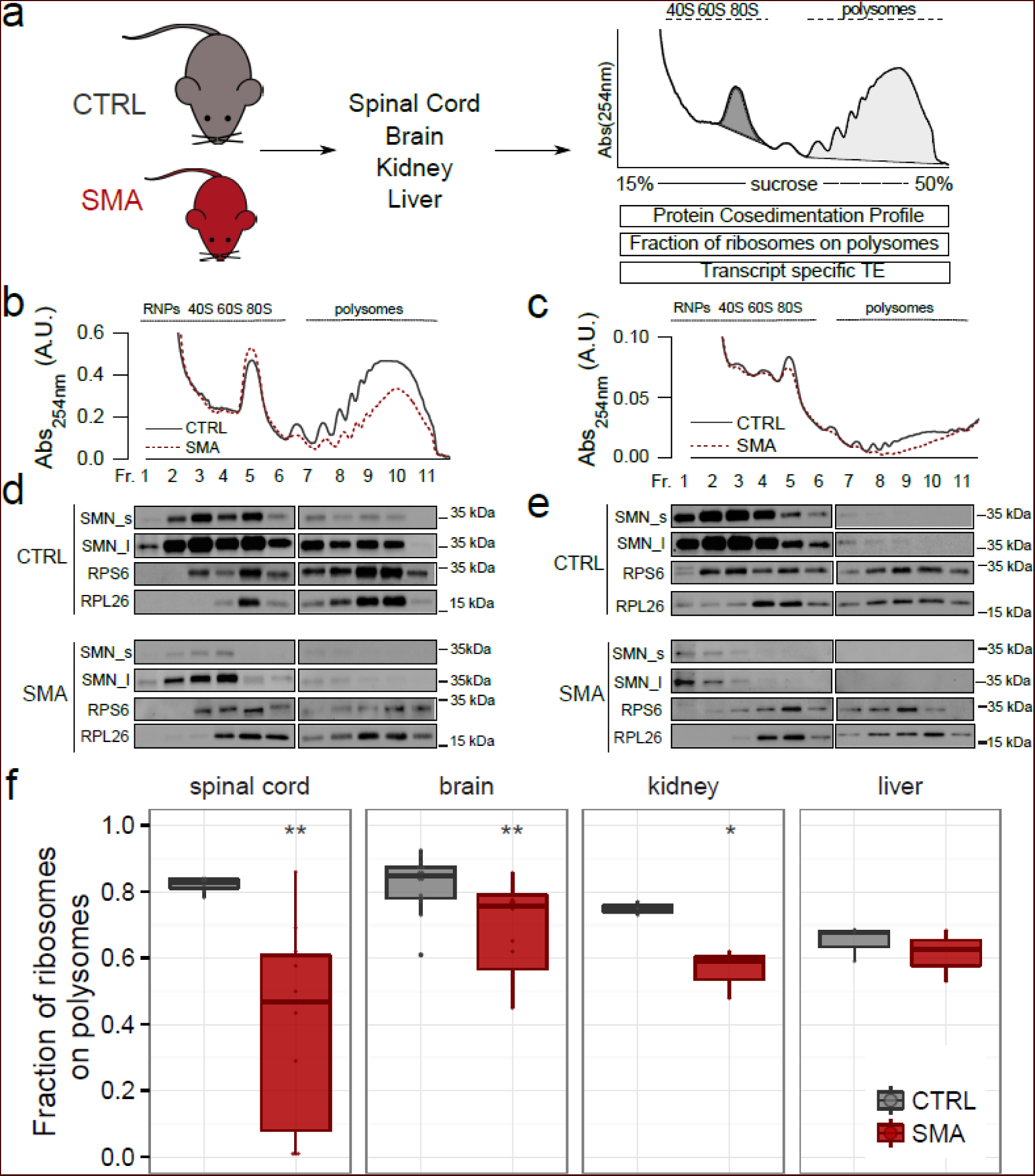
Translation is impaired in late-symptomatic SMA nervous tissues. (**a**) Experimental design used to obtain polysomal profiles from control (CTRL) and SMA mouse tissues. After purification of polysomes it is possible to; i) extract proteins from fractions and analyse co-sedimentation profiles with ribosomes and polysomes; ii) calculate the fraction of ribosomes on polysomes, as the ratio between the area under the curve of polysomes (light grey fill) and the area under the curve of polysomes plus the area of the 80S peak (dark grey fill); iii) extract the polysomal RNA and the total cytoplasmic RNA and analyse the translation efficiency of associated transcripts by Next Generation Sequencing. (**b,c**) Sucrose gradient absorbance profiles from CTRL and SMA brains and spinal cords (late symptomatic). The first peak contains free cytosolic light components (RNPs), the following peaks include ribosomal subunits (40S and 60S) and not translating monosomes (80S). The remaining peaks of the profile represent polysomes. (**d,e**) Co-sedimentation profiles of SMN, RPS6 and RPL26 under the corresponding sucrose gradient fractions by western blotting. To appreciate the distribution of SMN along the profile, the membranes were developed using both short (SMN_s) and long (SMN_l) exposure times. (**f**) Comparison between the fraction of ribosomes in polysomes in CTRL and SMA mouse tissues at late symptomatic stage (spinal cord: CTRL n=7, SMA n=10; brain: CTRL n=16, SMA n=14; kidney: CTRL n=3, SMA n=3; liver: CTRL n=3, SMA n=3, * P<0.05, ** P<0.01, *** P<0.001, two-tailed t-test No significant differences were observed in the liver of SMA mice compared to controls.

The cytoplasmic localization of SMN, alongside its known role in mRNA transport along the axon and its potential association with both ribosomal proteins [44] and polysomes [29, 33], prompted us to confirm its association with polysomes *in vivo*. To do that, we determined the co-sedimentation profiles of SMN in control and SMA brains and spinal cords by immunoblotting (Fig. 1d and e). The ribosomal proteins RPS6 and RPL26 were used as markers of the 40S and 60S subunits of the ribosome, respectively. We observed co-sedimentation of SMN with the ribosomal subunits, the monosomes (80S) and polysomes in both brain (Fig. 1d) and spinal cord (Fig. 1e) from control mice. Co-sedimentation profiles generated from SMA mice demonstrated that, as expected, SMN protein abundance was strongly reduced and SMN co-sedimentation with polysomes was lost (Fig. 1d and e). To further demonstrate the association of SMN with polysomes, we treated the lysates with EDTA, to dissociate ribosomes into the small and large subunits of the ribosome. As expected, under these conditions, the sedimentation profile of SMN moved towards lighter fractions (**Supplementary Figure 1**). To exclude the association of SMN with hnRNPs and to further demonstrate the robust association of SMN with polysomes, we compared the co-sedimentation profile of hnRNP A1 to that of SMN under control and EDTA conditions, and found no co-sedimentation of hnRNP A1 with SMN (**Supplementary Figure 1**). Thus, SMN protein co-sediments with the translational machinery *in vivo*.

When comparing polysomal profiles from late symptomatic SMA and age-matched control tissue (Fig. 1b and c, see also **Supplementary Figure 2** for pre- and early symptomatic stages and **Supplementary Figure 3** for other tissues) we noted a reduction in the polysome peak of SMA profiles, suggestive of translational defects. To quantify this effect, we determined the fraction of ribosomes engaged on polysomes (FRP), calculated as the ratio between the absorbance of polysomes and the total absorbance of non-translating 80S ribosomes and polysomes (see: Materials and methods, Fig. 1f). This value provides an estimate of the translational status/activity of tissues and cells and describes the engagement of ribosomes on RNAs in polysomes and/or the recruitment of mRNAs on polysomes for translation. [31, 38, 45, 46]. Comparing this parameter in different tissues from late-symptomatic SMA mice and controls, we observed the strongest decrease in SMA spinal cord, modest decreases in brain and kidney, but no change in liver, suggesting that translational defects display tissue-specificity, with the most pathologically affected tissues in SMA showing the greatest defects.

### Translational impairment correlates with SMA disease progression

One major challenge in studying neurodegenerative diseases such as SMA is to distinguish true pathogenic changes, occurring early in pathogenesis, from downstream cellular phenotypes that represent later-stage consequences of the disease [47]. We therefore wanted to establish whether the translational defects we observed at late symptomatic stages were also an early hallmark of the disease. To this end, we calculated the fraction of ribosomes on polysomes (FRP) at pre- and early-symptomatic stage of the disease (Fig.2a and b). Although no significant changes were apparent at pre-symptomatic stages, we identified a significant decrease in FRP at early-symptomatic stages in SMA brain, indicating that translation can be affected early in the disease process (Fig. 2a; early symptomatic, control: FRP=0.85±0.02, SMA: FRP=0.71±0.07, P=3.6E-03).

**Figure 2.**
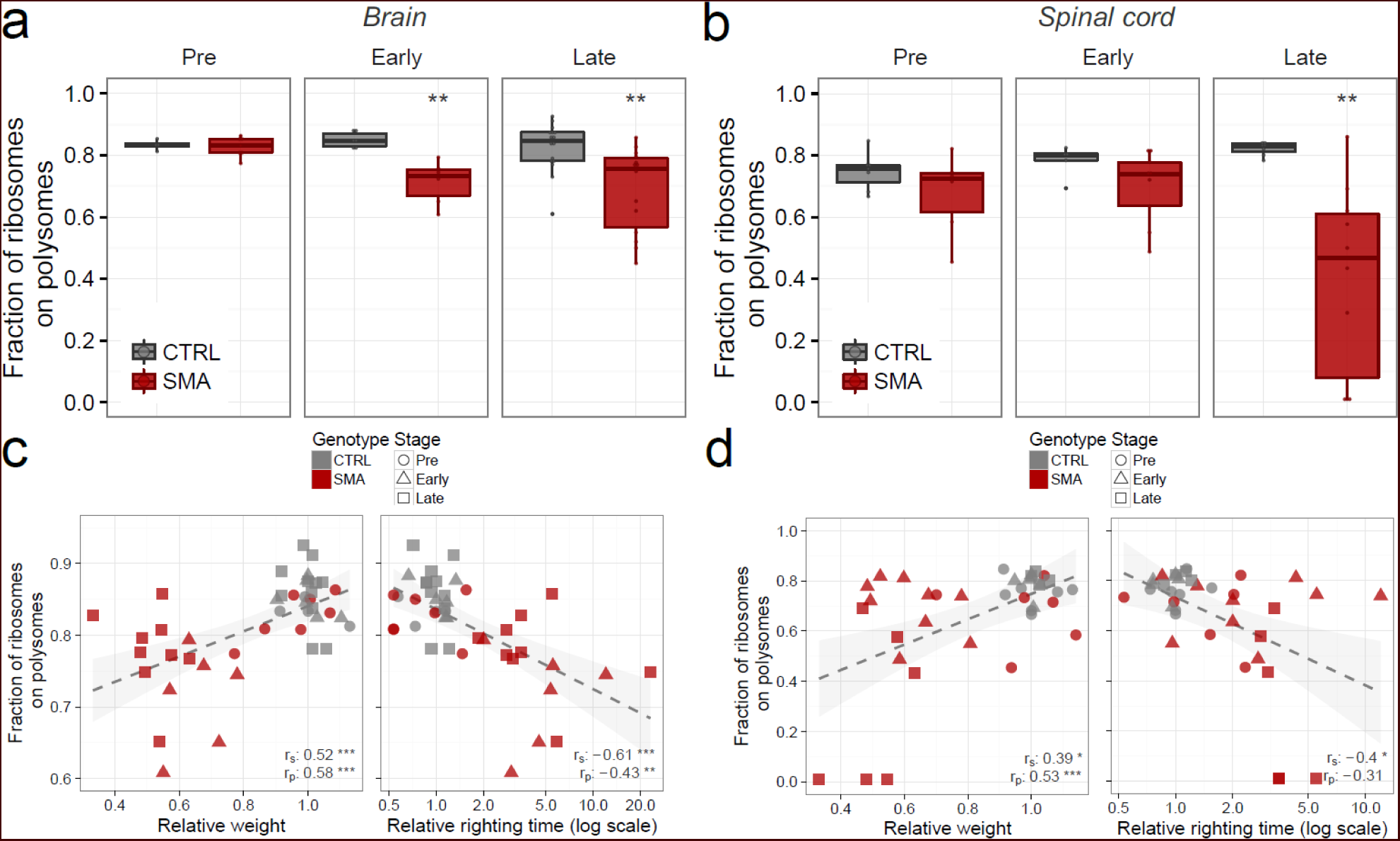
Translational impairment occurs early and correlates with disease progression in SMA. (a,b) Comparison between the fraction of ribosomes in polysomes in CTRL and SMA mouse brains (a) and spinal cords (b) at three stages of disease (brain: pre-symptomatic: ctrl n=4, SMA n=7; early symptomatic: ctrl n=6, SMA n=6; late symptomatic: ctrl n=16, SMA n=14; spinal cord: pre-symptomatic: ctrl n=7, SMA n=6; early symptomatic: ctrl n=5, SMA n=9; late symptomatic: ctrl n=7, SMA n=10, * P<0.05, ** P<0.01, *** P<0.001, two-tailed t-test). **(c,d)** Relationship between body weight (left panels) or righting time (right panels) and the corresponding fraction of ribosomes on polysomes, obtained from CTRL and SMA mouse brains (c) and spinal cords (d). Each point corresponds to data from one individual mouse. Spearman and Pearson correlations between the two measures displayed in the scatterplot are indicated (* P<0.05, ** P<0.01, *** P<0.001, correlation test).

To investigate the temporal nature of translational changes in more detail, we compared the extent of translational impairment with established SMA phenotypic disease readouts: body weight [37] and righting reflex time [48]. In agreement with previous studies, SMA mice were easily discernible from their control littermates at early- and late-symptomatic stages, as demonstrated by decreased body weight and increased righting time (**Additional file 2**). In contrast, at pre-symptomatic stages, SMA and control mice were indistinguishable. We next correlated body weight and righting time with the corresponding fraction of ribosomes on polysomes obtained from each individual mouse (Fig. 2c and d). Control and pre-symptomatic SMA mice showed a robust co-occurrence of high body weight and short righting time with a high FRP in brain (Fig. 2c) and spinal cord (Fig. 2d). In contrast, results obtained from both early- and late symptomatic SMA mice were clearly distinct (**Supplementary Figure 4**), and a progressive worsening of disease symptoms accompanied a parallel decline in FRP in both the spinal cord and the brain. Correlation analyses for relative weight vs FRP (Fig. 2c (brain) and d (spinal cord), left panels) and righting time vs FRP (Fig. 2c (brain) and d (spinal cord), right panels) confirmed that translational defects in SMA are associated with established phenotypic read-outs of disease progression. Taken together, these results show that a decrease in FRP is an early event during SMA pathogenesis, with defects becoming progressively more severe as the disease progresses.

### Restoration of SMN protein in late symptomatic SMA mice shows that translational defects are SMN-dependent

Having shown that the fraction of ribosomes on polysomes was decreased in SMA, and that translational impairment correlates with disease severity, we next wanted to establish whether these changes were directly dependent upon SMN protein levels. To assess this, we investigated the translational activity by determining the FRP in SMA mice that had been treated with an established ASO that leads to restoration of SMN levels [49, 50]. SMA mice were treated with a single intravenous injection of a 25-mer morpholino ASO (PMO25) that targets the *SMN2* intron 7 splicing silencer N1. Following treatment, *SMN2* exon 7 inclusion in the central nervous system (CNS) increased over two-fold [50]. Exon 7 inclusion leads to increased SMN protein levels (**Supplementary Figure 5**) and robust rescue of symptoms compared with non-treated SMA mice, as illustrated by increased body weights and decreased righting times (**Supplementary Figure 6**), as well as increased survival [50]. We performed polysomal profiling after ASO treatment (Fig. 3a) and determined the FRP by analysing polysomal profiles from brain and spinal cord of control, SMA and ASO-treated SMA mice (Fig. 3b and Fig. 3c). Loss of SMN from polysomes in late symptomatic SMA mice was associated with a significantly decreased FRP in brain and spinal cord compared to littermate controls (Fig. 3c, brain control: FRP=0.75±0.09; SMA: FRP=0.53±0.06, P=1.1E-03; spinal cord control: FRP=0.84±0.01; SMA: FRP=0.57±0.24, P=5.4E-02). This effect was completely rescued by ASO treatment which restored both the amount of SMN on polysomes and the fraction of ribosomes in polysomes (Fig. 3c, ASO: FRP=0.86±0.02, P=8.4E-05; ASO: FRP=0.77±0.016, P=1.17E-01).

**Figure 3.**
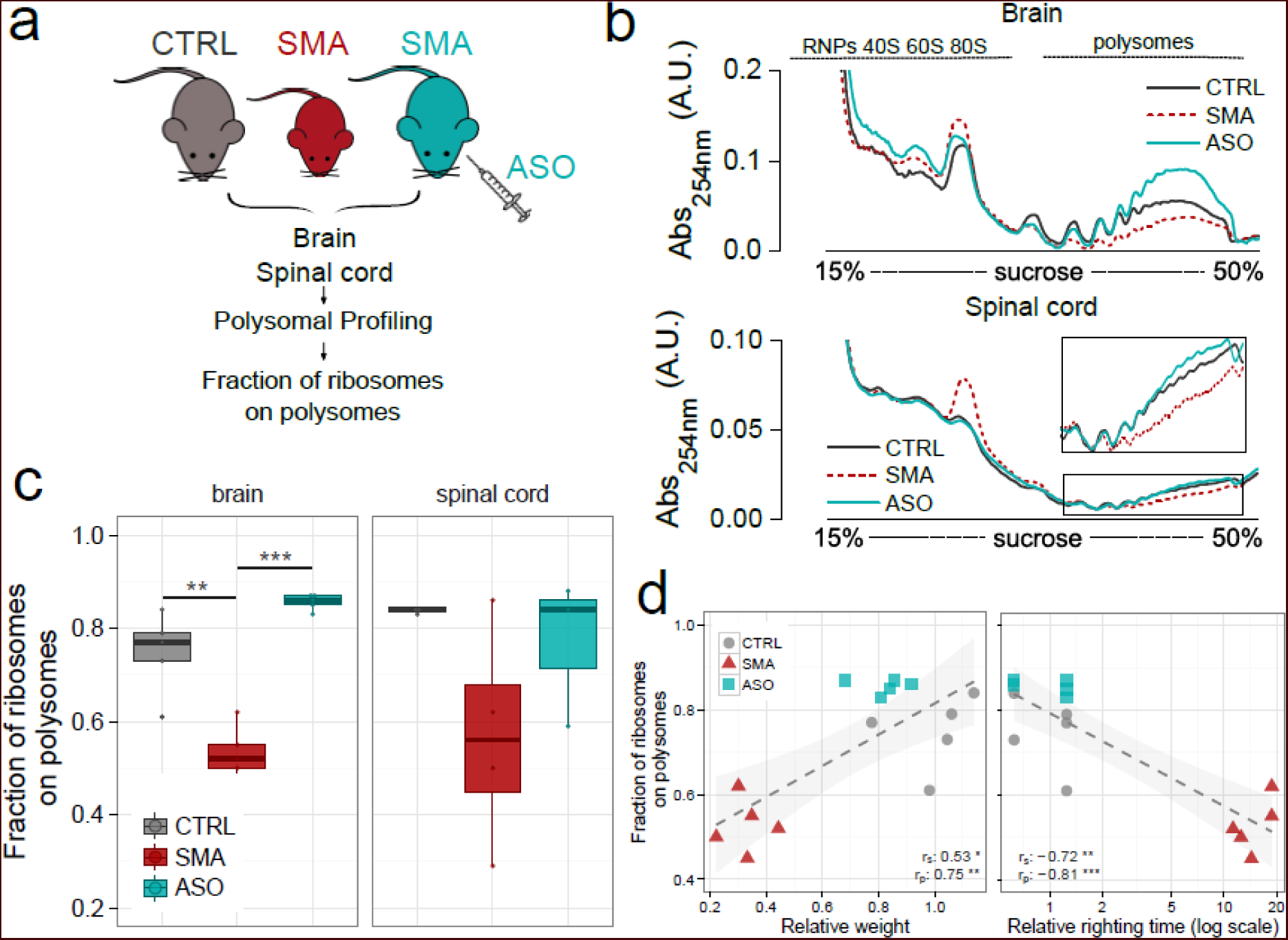
Translational defects are rescued by treatment with an ASO restoring SMN. (**a**) Experimental design: polysomal profiling was performed on brains from CTRL mice, late symptomatic SMA mice, and late-symptomatic SMA mice treated with antisense oligonucleotide PMO25 (ASO). Polysomal profiles were generated and the fraction of ribosomes on polysomes was calculated as described in Figure 1. (**b**) Representative sucrose gradient absorbance profiles obtained from brains (upper panel) and spinal cords (lower panel) in each experimental group. (**c**) Box-whisker plots of the fraction of ribosomes on polysomes from CTRL, late symptomatic SMA and SMA-ASO brains (left panel) and spinal cords (right panel) (CTRL, n=5; SMA, n=5; ASO, n=5, * P<0.05, ** P<0.01, *** P<0.001, one-tailed t-test). (**d**) Relationship between body weight (left panel) or righting time (right panel) and the corresponding fraction of ribosomes on polysomes, obtained from CTRL, SMA and ASO-treated mouse brains. Each point corresponds to data from one individual mouse. Spearman and Pearson correlations between the two measures displayed in the scatterplot are indicated. (* P<0.05, ** P<0.01, *** P<0.001, correlation test).

Subsequent correlation of body weight and righting time with FRP confirmed the robust rescue as ASO-treated mice were indistinguishable from control mice (Fig. 3d and **Supplementary Figure 7**). Taken together, these results demonstrate that restoring SMN expression to normal levels rescues impairments in translation. Thus, translational defects observed in SMA are reversible and depend on SMN expression.

### Translational impairment is most pronounced in primary affected brain regions and cell types in vivo and in vitro

The findings described above demonstrate that defects in ribosomal engagement on polysomes are a pronounced feature of SMA pathogenesis. However, these translational defects had only been observed at the level of whole tissues (e.g. spinal cord or brain) whilst pathological changes occurring in SMA are known to be restricted to specific anatomical regions and cell populations (e.g. motor neurons) within the brain and spinal cord [23]. We therefore wanted to investigate whether translational changes were more pronounced in particularly affected brain regions from SMA mice. First, we dissected three brain regions from control and late-symptomatic SMA mice and analysed them by determining the FRP by polysome profiling (Fig. 4a). In line with previously observed pathological changes [51], we found significant defects in the cortex and hippocampus (both regions known to be affected in mice modelling severe SMA), while no changes were detected in the cerebellum (a region largely unaffected in SMA mice).

**Figure 4.**
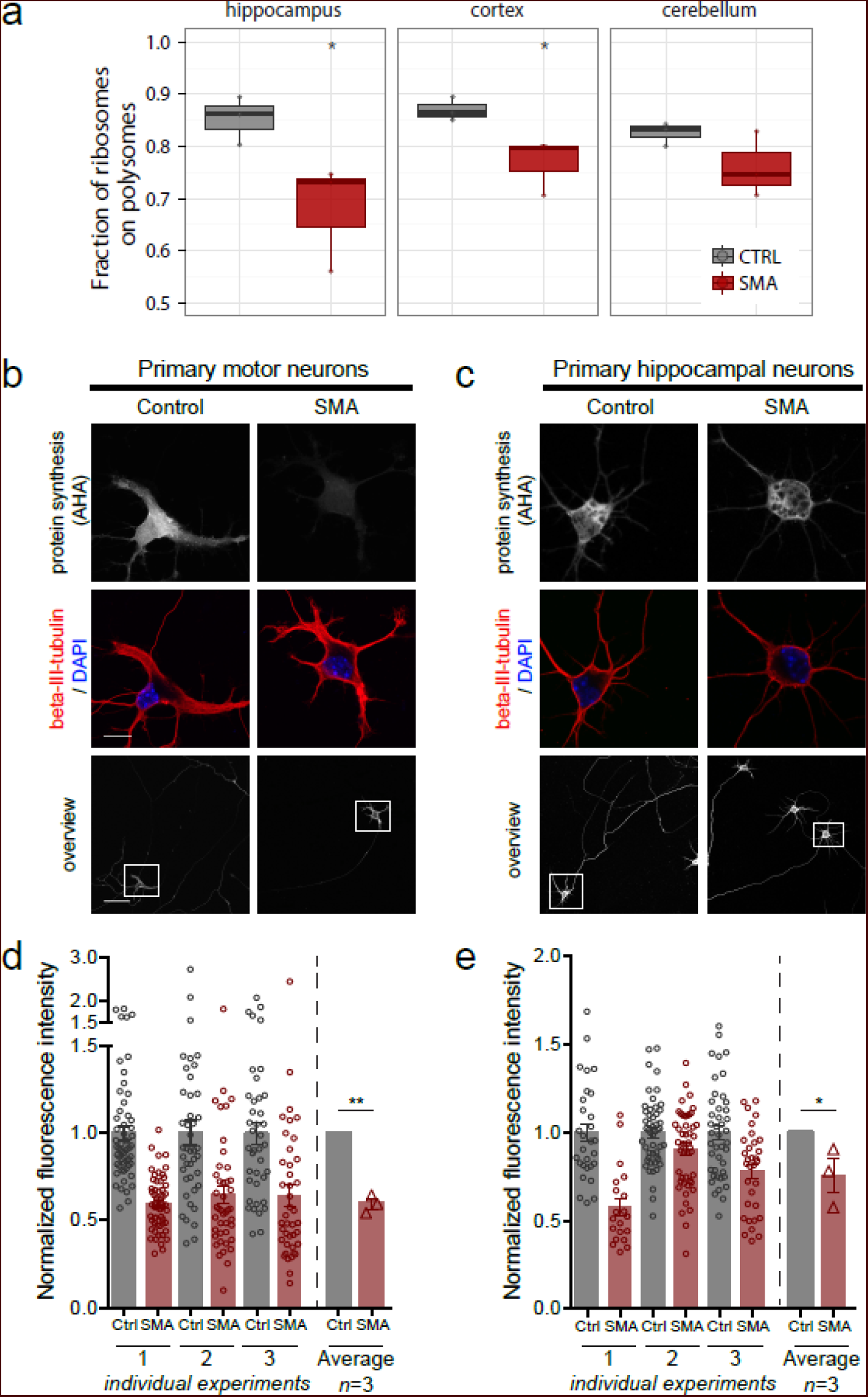
Translation defect is cell autonomous and dependent on SMN loss. (**a**) Comparison between the fraction of ribosomes on polysomes in differentially affected brain regions from CTRL and SMA mouse brain regions: hippocampus (most affected), cortex and cerebellum (least affected) were analysed in late symptomatic mice (for each region 3 Ctrl and 3 SMA samples were analyzed). Significant reductions between CTRL and SMA brain regions were identified with one-tailed t-test (*P<0.05). (**b,c**) Primary motor neurons (**b**) and primary hippocampal neurons (**c**) from control and SMA embryos were stained to reveal overall morphology (beta-III-tubulin, red) and nuclear integrity (DAPI, blue). Protein synthesis was visualized by labelling newly synthesised proteins using L-azidohomoalanine (AHA, grey scale). Beta-III-tubulin staining was used to assess overall neuronal morphology (bottom panel). (**d**) AHA fluorescence intensity of primary motor neurons was normalized to control and determined for individual SMA (n=43, 41, 56 for each of the respective experiments) and control (n=42, 40, 56 for each of the respective experiments) neurons in each of 3 independent primary neuron preparations; the final graph shows the mean value obtained from the 3 experiments. (**e**) AHA fluorescence intensity of primary hippocampal neurons was normalized to control and determined for individual SMA (n=29, 50, 48 for each of the respective experiments) and control (n=20, 48, 32 for each of the respective experiments) neurons in each of 3 independent primary neuron preparations; the final graph shows the mean value obtained from the 3 experiments. (Scale bars: 50 µm (overview), 10 µm (cell bodies; beta-III-tubulin/DAPI and AHA. DAPI, 4’,6-diamindion-2-phenylindole; **P< 0.01, *P<0.05, student’s t-test; error bars ± SEM.)

To determine whether the decrease in FRP was both cell autonomous and dependent on SMN loss, we next developed motor-neuron like cell lines (NSC-34) expressing SMN at different levels using CRISPR/Cas9 technology (see **Supplementary Methods**). We isolated and expanded two clones expressing 20% and 0% SMN, respectively (**Supplementary Figure 8a**) and performed polysome profiling (**Supplementary Figure 8b**) to determine the FRP for each of the cell lines. Here, we observed similar translational defects as seen in tissues that, moreover, correlated with the level of SMN loss (**Supplementary Figure 8c**).

Next, we cultured primary motor neurons (the most affected neuronal cell type [52]) and primary hippocampal neurons (a more modestly affected neuronal cell type [51]) from SMA and control mouse embryos. We measured *de novo* protein synthesis levels using an established metabolic labelling technique, based on incorporating the methionine-homologue L-azidohomoalanine (AHA) into newly synthesized proteins [53]. Imaging the relative fluorescence of AHA-labeled proteins provides an indication of the translational activity of cultured cells. When cultured primary motor neurons were treated with anisomycin, an inhibitor of translation, the AHA fluorescence signal was reduced to background levels (**Supplementary Figure 9**), indicating that this assay provides a specific signal when used on primary motor neurons. When applying AHA metabolic labelling to primary SMA-derived motor and hippocampal neuron cultures, we observed a marked decrease in fluorescence intensity compared to primary cultures derived from control embryos (Fig. 4b and fig. 4c). Quantification of this finding across 3 independent primary motor neuron preparations revealed a 37% decrease in fluorescence intensity when comparing SMA to control motor neurons (Fig. 4d). Comparable quantification in primary hippocampal neurons indicated a more modest decrease in *de novo* protein synthesis of 21% (Fig. 4e). These results confirm that the decreases in FRP previously observed at the level of whole tissues are compatible with defects occurring at the level of individual neurons, with the most robust defects in translational observed in the primary affected cell type in SMA, the motor neuron.

Taken together, the findings that *de novo* protein synthesis is impaired at the level of individual neurons and that FRP decrease is both brain region specific and correlates with SMN levels, suggest that the translational deficit is acell-autonomous event that depends specifically on the loss of SMN expression.

### POL-Seq analysis reveals genome-wide impairment of translation efficiencies in early- and late-symptomatic SMA mice

Having identified robust translational defects in tissues, cell lines and individual primary motor neurons, we next went on to establish the consequences of these defects in the recruitment of mRNAs onto polysomes in SMA. We employed next generation sequencing (NGS) to identify and quantify, in parallel, the total cytoplasmic RNA (RNA-Seq) and the RNAs associated with polysomes (POL-Seq). This approach enables the simultaneous identification of variations in RNA populations at both *translatome* (RNAs engaged with polysomes) and *transcriptome* (steady state cytoplasmic RNAs) levels.

We extracted both total cytoplasmic and polysomal RNAs from pooled sucrose fractions from control, early- and late-symptomatic SMA mice, as illustrated in Fig. 5a. As these experiments represent the first application of translatome profiling to SMA tissues during disease progression, we included the complete POL-Seq and RNA-Seq datasets in a fully annotated, searchable and filterable format (**Additional File 3**). Among the most up and down-regulated transcripts in SMA (Fig. 5b), we chose a set of nine differentially expressed genes that could be reliably detected using qPCR at both early and late symptomatic stages for subsequent validation. We confirmed the reproducibility of the NGS data by qPCR (**Supplementary Figure 10**), finding a high concordance between the two techniques at both stages (r^2^ =0.77 for early symptomatic stage, r^2^ =0.99 for late symptomatic stage).

**Figure 5.**
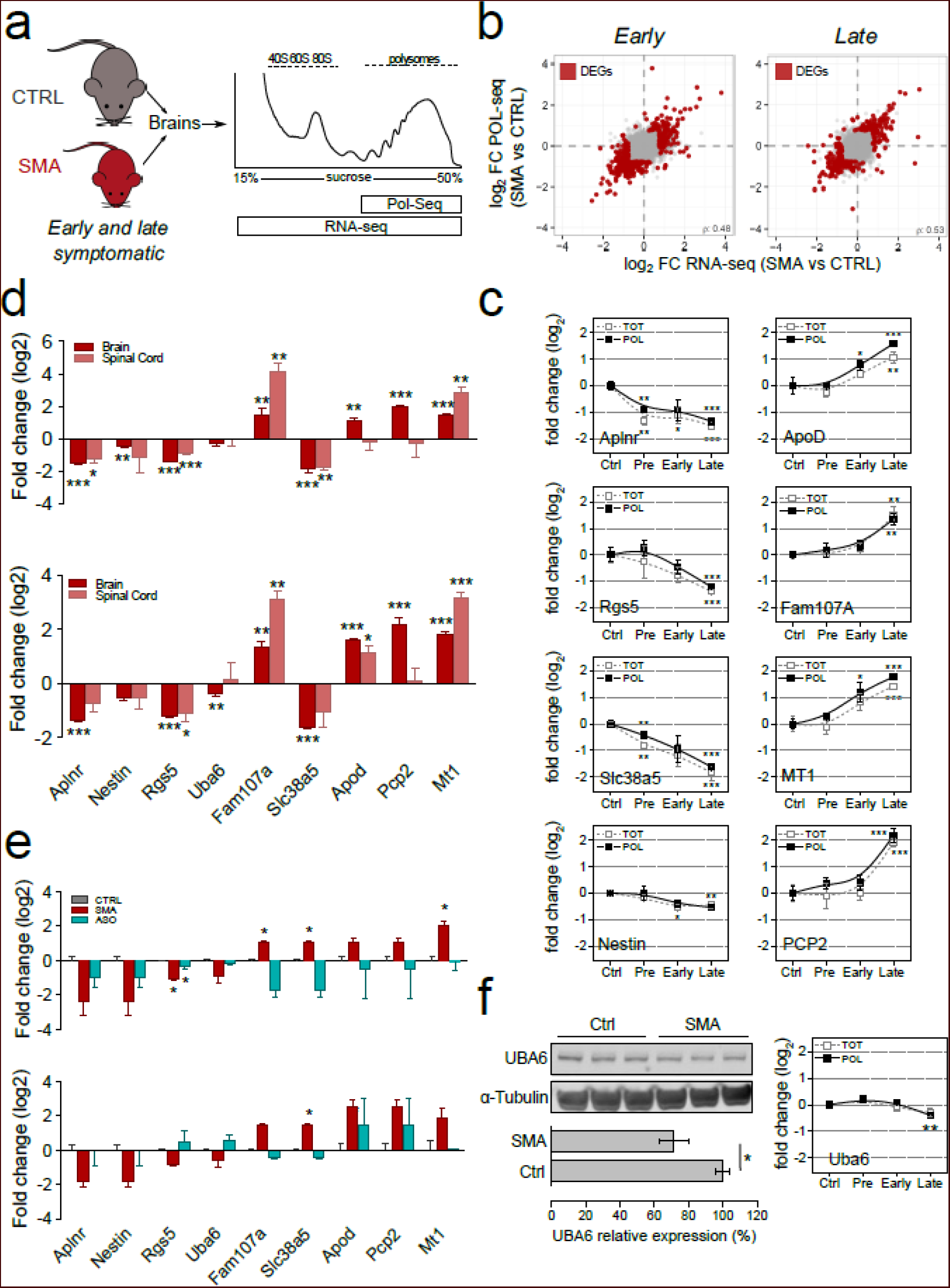
Transcriptome and translatome profiling reveal gene expression alterations in SMA mice. (**a**) Experimental design used to perform POLseq and RNAseq profiling on brains from late symptomatic SMA mice and controls. (**b**) Scatterplots displaying, for each quantified transcript, translatome (POL-seq, y-axis) and transcriptome (RNA-seq, x-axis) variations between SMA and control mice. Differentially expressed genes (DEGs) are labelled in red. (**c**) Time-course of changes in levels of differentially expressed genes across three stages of disease with respect to the control brains at the transcriptional (TOT) and translational levels (POL). Analyses were performed at pre-, early and late symptomatic stages for DEGs identified in late symptomatic tissue. All samples were normalized to the geometric mean value of actin and cyclophilin a. For each transcript the mean value ± SEM is shown (2-3 biological replicates and 2-4 technical replicates; * P<0.05, ** P<0.01, *** P<0.001, two-tailed unpaired t-test). (**d**) Comparison of differentially expressed genes between tissues (brain and spinal cord, late symptomatic) at transcriptional (upper panel) and translational levels (lower panel). All samples were normalized to the geometric mean value of actin and cyclophilin a. For each transcript the mean value ± SEM is shown (2-3 biological replicates and 2-4 technical replicates; * P<0.05, ** P<0.01, *** P<0.001, two-tailed unpaired t-test). (**e**) Comparison of differentially expressed genes after SMN-targeted treatment (Ctrl, SMA and ASO-treated brain, late symptomatic) at transcriptional (upper panel) and translational levels (lower panel). All samples were normalized to the geometric mean value of actin and cyclophilin a. For each transcript, the mean value ± SEM is shown (2 biological replicates and 4 technical replicates; * P<0.05, ** P<0.01, *** P<0.001, two-tailed unpaired t-test). (**f**) Consistency in differential expression of Uba6 in SMA compared to control at RNA (right panel) and protein level (left panel). Right panel: time-course of changes in levels of Uba6 across three stages of disease with respect to the control brain at the transcriptional (TOT) and translational levels (POL). The mean value ± SEM is shown (2 biological replicates and 4 technical replicates; ** P<0.01, two-tailed unpaired t-test). Lower panel: protein expression levels of Uba6 in whole brain from control and SMA mice. Alpha-tubulin was used as a loading control (n=6; * P<0.05, two-tailed unpaired t-test; error bars ± SEM).

To understand if the variations in mRNA we previously identified at early- and late-symptomatic stages were also hallmarks of pre-symptomatic stages of the disease, we determined an expression time-course for the same transcripts that were selected for qPCR validation (Fig. 5c). This showed that, although most transcripts were unchanged at the pre-symptomatic stage, several transcripts were differentially expressed, indicating some early pathogenic changes. Next, as our initial NGS and validation experiments were carried out on RNA isolated from brain tissue, we went on to determine the expression levels of our validation panel of transcripts in spinal cord. Importantly, the majority of changes observed at both early and late-symptomatic stages could be detected not only in brain but also in spinal cord (Fig. 5d). These findings suggest that the changes captured at symptomatic stages are *bona-fide* features of subtle defects occurring at early stages of disease progression in disease-affected tissues.

Importantly, these changes were reversible, as ASO treatment (see Fig. 3) normalised the expression at both transcriptional and translational levels (Fig. 5e). To confirm that at least some of the changes identified in SMA at the polysomal level resulted in corresponding changes at the protein level, we measured levels of UBA6 in SMA mice. This transcript was down-regulated only at the translational level and showed a concomitant decrease in protein abundance, confirming the accuracy of our translation measurements (Fig. 5f).

Next, to establish the global effect of impaired translation in SMA on individual RNA transcripts, we used the ratio between POL-Seq and RNA-Seq abundances to determine and compare the translation efficiencies (TE) of individual RNA transcripts in control and SMA tissues [54, 55]. This analysis allowed us to highlight transcripts that have a significantly altered TE between control and SMA at the transcriptome-wide level, both at early and late symptomatic stages of the disease (Fig. 6a). We found that 470 and 442 transcripts were significantly altered in TE at the early and late symptomatic stages, respectively (**Supplementary Data 2**). Interestingly, most RNAs with altered TE were protein coding, but a substantial proportion of these RNAs were noncoding (long noncoding and antisense being the most represented categories, Fig. 6b).

**Figure 6.**
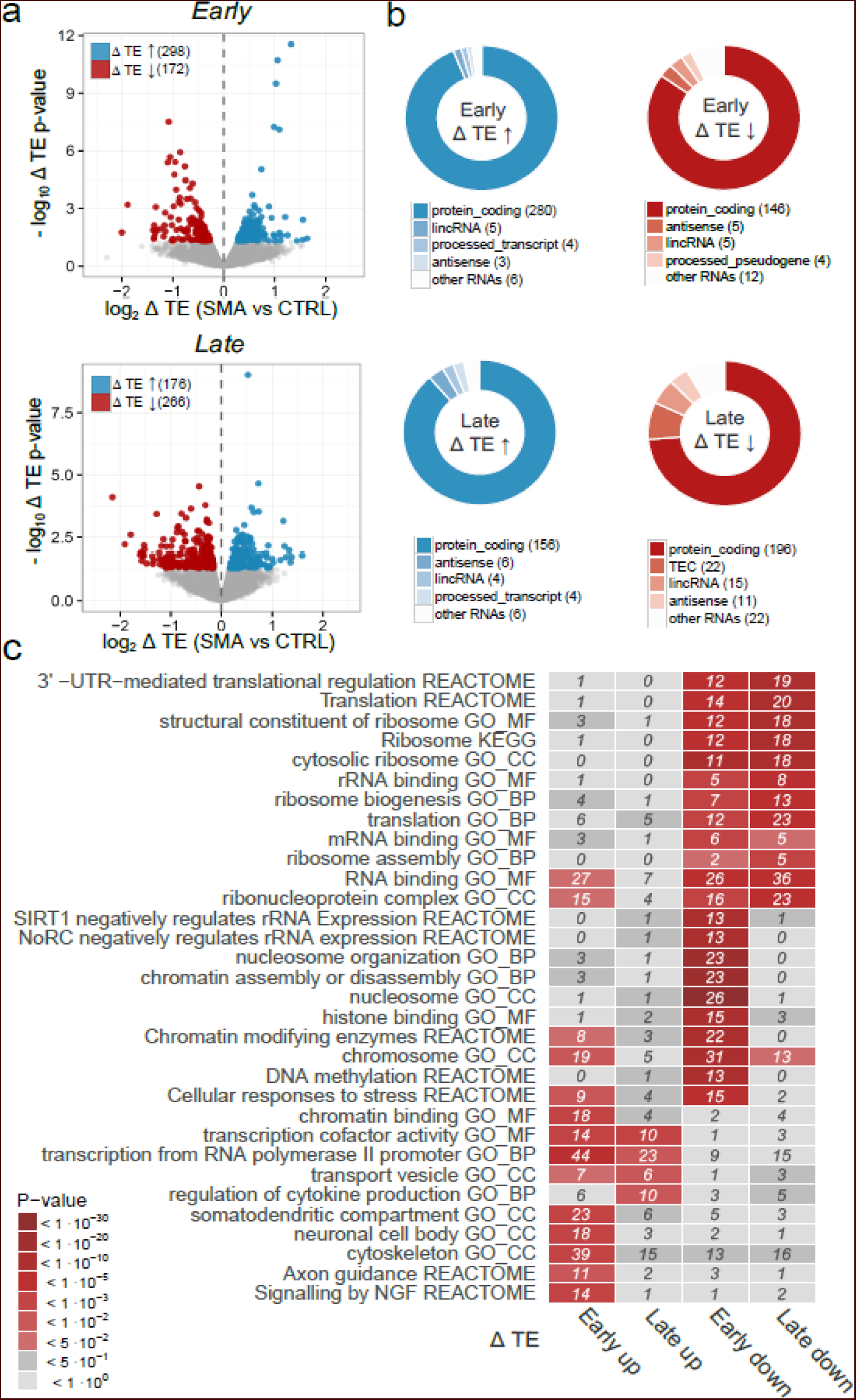
Translation Efficiency analysis identifies functional clusters of RNAs with altered translation in SMA. (**a**) Volcano plot displaying translation efficiency variations (x-axis) and associated p-values (y-axis) in early symptomatic (upper panel) and late symptomatic (lower panel) SMA mouse brain compared to controls. TE analysis was performed with Xtail (see Methods). Genes with statistically significant variations in TE are colour labelled according to the direction of the observed change (up (blue) or down (red) regulation in SMA). (**b**) RNA subtypes of genes with significantly altered TE in early- (upper panels) and late-symptomatic (lower panels) SMA mice. (**c**) Heatmap showing and comparing top enriched terms (from Gene Ontology) and pathways (from KEGG and Reactome). Enrichment analysis was performed on the lists of genes with significant changes of TE in early- and late-symptomatic SMA brains. Significant enrichments are displayed in red shades. The number of genes contributing to the enrichment is indicated in each tile.

To better understand the molecular processes associated with these mRNAs, we performed Gene Ontology and pathway enrichment analyses specifically on transcripts that had changed TE values in SMA at either early or late stages of the disease (**Additional File 4**). Focusing our attention initially on transcripts with reduced levels of TE at early stages of disease, we found strong enrichments in two major cellular processes: nucleosome/chromatin and ribosome/translation. Interestingly, however, when examining transcripts with decreased TE, only the ribosome/translation category was robustly conserved between early and late stages of disease (Fig. 6c). This suggests that defects in ribosome/translation pathways are an early event in SMA pathogenesis, persisting into late stages of the disease.

### Defects in ribosome biology are hallmarks of SMA

After having established that genome-wide changes in TE underlie defects in ribosome/translation occurring during SMA pathogenesis, we identified transcripts with a significant reduction of TE and associated with the Reactome pathway “translation” (Fig. 6c), revealing a large majority of ribosomal proteins, with factors involved in translation initiation and elongation (Fig. 7a). To validate these findings, we quantified individual transcript levels for fourteen ribosomal genes and one translation factor by qPCR (Fig. 7b). These experiments confirmed that the expression levels of numerous individual genes involved in ribosome biogenesis [56–58] are affected in SMA, both at early- and late-symptomatic stages of disease and suggest that defects in ribosome abundance could underlie, at least in part, the translational defects that occur in SMA.

**Figure 7.**
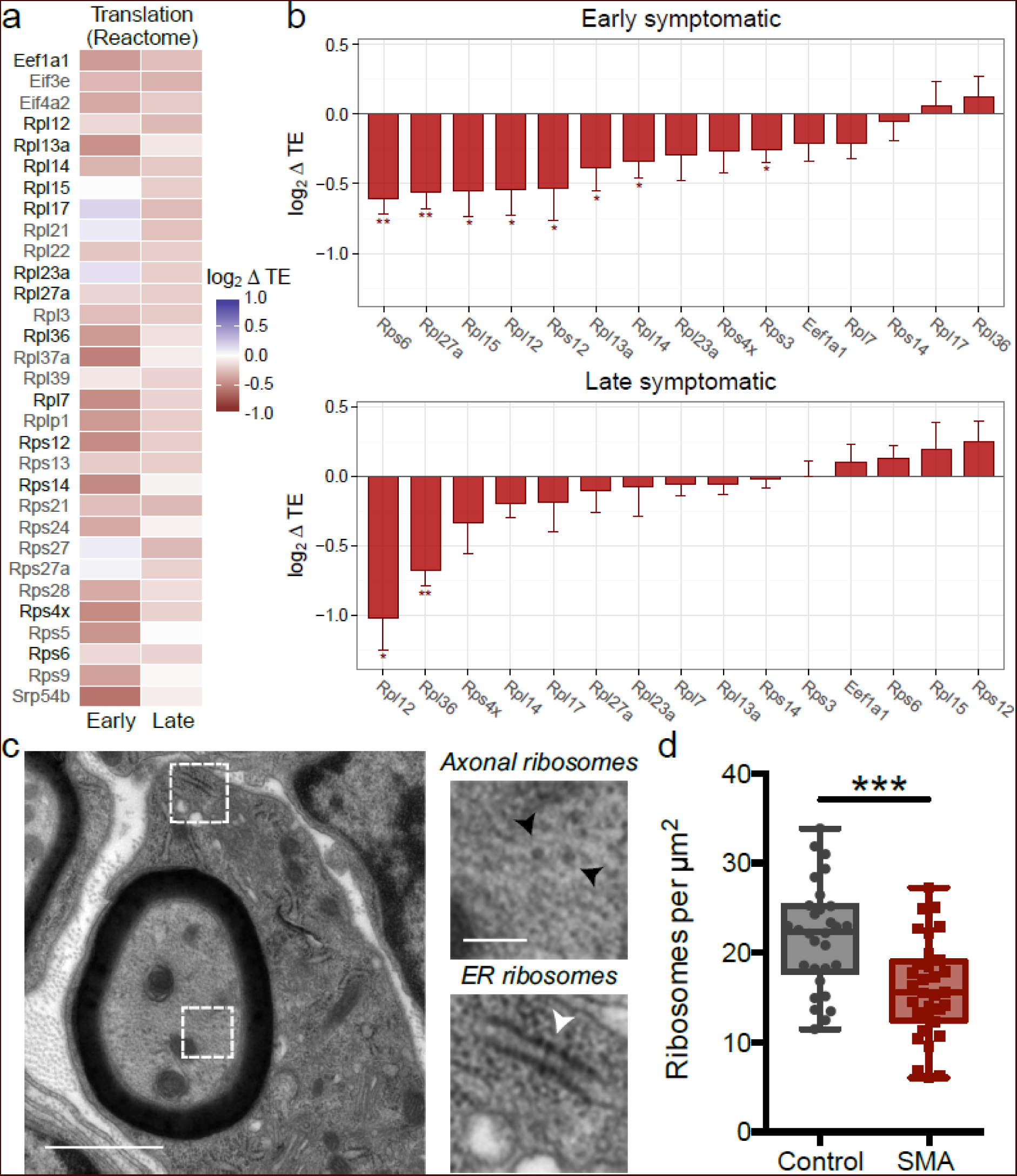
Translational alterations in SMA are associated with defects in ribosomes. (**a**) Heatmat displaying all genes with altered translational efficiency (TE) in SMA brains and annotated under the “Translation” Reactome pathway, significantly enriched in Fig. 6c. The majority of genes belong to the family of ribosomal proteins. Genes further analysed by qPCR are highlighted in black. (**b**) qPCR derived variations of translation efficiency for ribosomal proteins and one elongation factor from (**a**) in early (upper panel) and late (lower panel) symptomatic mouse brains. (Mean value ± SEM is shown; 3-4 biological replicates and 2-6 technical replicates; all genes were normalized to the geometric mean value of actin and cyclophilin a; one-tailed unpaired t-test; * P<0.05, ** P<0.01, *** P<0.001). (**c**) Representative electron micrograph of a large diameter (motor) axon in the intercostal nerve from a P5 ‘severe’ SMA mouse (black arrowheads: axonal ribosomes, white arrowhead = ER ribosomes). (**d**) Counts of axonal ribosomes revealed a decrease in the density of ribosomes in SMA mouse axons compared to controls (n=30 axon profiles; N=3 mice per genotype; *** P<0.001, two-tailed unpaired t test).

To prove this hypothesis and investigate whether the down-regulation of translation-related transcripts was reflected by defects in ribosome abundance in motor neurons *in vivo*, we performed ultrastructural analyses of ribosomes in banked samples of intercostal nerves isolated from symptomatic severe SMA mice and littermate controls, where neuropathological changes have previously been established [59]. As expected [60], transmission electron microscopy revealed prominent axonal ribosomes in both control and SMA mice (Fig. 7c). Quantitative analyses of axonal ribosomal density revealed a 27% decrease in the number of axonal ribosomes in intercostal nerves from SMA mice versus controls (Fig. 7d). Thus, defects in translational pathways and ribosome biology that occur at the molecular level in SMA are accompanied by structural defects in the ribosomal machinery.

## Discussion

Increasing evidence suggests a role for translational defects in the pathogenesis of a variety of neurodegenerative diseases. Ribosomal defects have been documented in conditions such as Alzheimer’s disease, Huntington’s disease, Parkinson’s disease and frontotemporal dementia [5–7]. Furthermore, translation factors are mutated in Vanishing White Matter disease [61], in epilepsy and in mental retardation [62–64]. Here we found that the SMN protein, whose loss causes SMA, is an important mediator of translation biology in the nervous system. Depletion of SMN in SMA resulted in a range of translational defects in affected tissues and cells. We employed a range of methods to specify the role of SMN in regulating translation *in vitro* and *in vivo*, using next generation sequencing approaches to determine transcriptome-wide changes that occur in the translatome in SMA pathogenesis. The identification of a transcriptome-wide set of differentially translated genes in SMA provided an important advancement in our understanding of motor neuron disease biology, as it allowed the determination of specific changes that occur downstream of translational defects. We also established that SMN depletion leads to an early and robust impairment of translation-related transcripts in SMA, and consequent defects in ribosomal biology that could provide a promising biomarker candidate for SMN-targeted clinical trials for SMA. In addition, we demonstrated that translational defects are cell-autonomous and can be rescued by therapies restoring SMN protein levels.

Our findings provide the first *in vivo* evidence to support the hypothesized physiological role of SMN in the localization of the translation machinery and translation of specific transcripts proposed previously in *in vitro* studies [33–36]. In accordance with our findings, SMN has been shown to localize to the nucleolus, where it interacts with U3 snoRNA, which is involved in the processing of ribosomal RNA (rRNA) [26]. Similarly, the characterization of the SMN interactome identified likely interactions between the SMN complex and numerous ribosomal proteins, as well as the translation elongation factor eEF1A1 [44]. Moreover, another recent study identified a number of ribosomal proteins amongst the most prominently down-regulated transcripts and proteins at vulnerable neuromuscular junctions in SMA [65].

One longstanding issue in several neurodegenerative diseases is the lack of a robust explanation for the cell-type specificity in the presence of defects of ubiquitously expressed proteins. Indeed, this is also a hallmark of the SMA disease following ubiquitous depletion of SMN protein [66]. Although spinal motor neurons are the most prominently affected cell type, numerous other cell-types and tissues can also be affected in SMA, albeit to a lesser extent, both in patients and a range of animal models of the disease [67–69]. The translational defects we report here were notably most pronounced in spinal cord and brain, with more modest defects observed in the kidneys and no observable defects in the liver. These findings support a threshold model for the requirement of SMN, where different tissues and cell-types require different minimal levels of SMN [70]. Our data would therefore suggest that defects in translation and ribosome biology are major downstream mediators of the effects of SMN on different tissues in SMA. In line with this model, we found that when comparing different CNS tissues (cortex, hippocampus and cerebellum) and cell-types (primary hippocampal and motor neurons), the tissues and cell-types known to be most affected in SMA showed the biggest defects in translational activity.

Taken together with findings from other neurodegenerative conditions, we might hypothesize that translation may represent a common defective feature across a variety of motor neuron diseases mediated by mutations in proteins involved in RNA metabolism. Indeed, several recent studies of the adult-onset motor neuron disease amyotrophic lateral sclerosis (ALS) have pointed towards a role for translational defects in motor neuron degeneration. For example, the main pathological protein in ALS, TDP43, has been shown to regulate the translation of specific transcripts [71]. Also, ALS-associated protein aggregates have been shown to affect the function of proteins that play an important role in regulating translation, such as FMRP [72, 73]. Moreover, EM analysis of patient nerves indicated the mislocalisation of ribosomes in ALS patients [74]. It is interesting to note that many proteins that have been implicated in regulating translation and have been associated to neurodegenerative diseases are known to interact as part of bigger RNA-protein complexes [73, 75–77]. This complicates the process of unravelling causative from secondary changes in disease, as likely functional redundancy in these complexes makes it challenging to detail true pathogenic changes. By showing that defects in translation are a prominent feature of SMA, we provide new insights into these challenging questions and wonder if defects in ribosome turnover, that is known to be tissue-specific, may play a role in producing tissue-specific defects in neurodegenerative diseases caused by ubiquitously expressed proteins involved in RNA metabolism.

## Methods

### Animal models and ASO treatment

The ‘Taiwanese’ mouse model of severe SMA was used in this study [37, 78, 79]. All mice were on a congenic FVB background, with colonies at the University of Edinburgh and UCL established from breeding pairs originally purchased from Jackson Labs. ‘Taiwanese’ SMA mice (*Smn-/-; SMN2tg/0*) carry two copies of *SMN2* on one allele and are on a null murine *Smn* background. Phenotypically normal heterozygous (*Smn-/+;SMN2tg/0*) littermates were used as controls. Mice were maintained following a breeding strategy previously described [78]. For EM experiments, tissue that had previously been banked from a ‘severe’ SMA mouse colony was used, as described previously [80]. Litters were retrospectively genotyped using standard PCR protocols. All mice were housed within the animal care facilities in Edinburgh and London under standard SPF conditions. All animal procedures and breeding were performed in accordance with University of Edinburgh and UCL institutional guidelines and under appropriate project and personal licenses granted by the UK Home Office.

Antisense oligonucleotide treatment was performed as previously described, using PMO25 that targets *SMN2* intron 7 splicing silencer N1 [50]. It is known that the disease progression of transgenic mice can vary between laboratories, even when the same transgenic model is used [81]. For clarity, throughout the paper we refer to the time points at which tissue was collected as pre-, early and late symptomatic throughout the paper. In Edinburgh, pre-symptomatic was P3, early symptomatic was P5 and late symptomatic was P7 (for figures 1-5 and S1-S5). The tissue obtained from UCL (figure 6 and S7-S9) was from late symptomatic (P10) mice that were phenotypically similar to P7 mice maintained in Edinburgh.

### Motor performance test

All mice used in the study were weighed daily. Motor performance was recorded using the righting reflex test, which is a common assay for neuromuscular function in neonatal mice [23, 82] and involves placing the mouse on its back on a flat surface and recording the time it takes to turn over onto its paws. If a mouse took longer than 30 seconds to right itself then the test was terminated. The righting test was performed in triplicate for each mouse and an average righting time was calculated.

### Tissue isolation

Neonatal mice were euthanized by overdose of anaesthetic, before decapitation to allow the brain to be removed. Spinal cord dissection was performed by first removing the spinal column and then using a needle and syringe to flush out the spinal cord with phosphate buffered saline (PBS) [52]. This technique leads to the isolation of intact spinal cord, which is important for sample preparation for polysome analysis. Peripheral tissues (kidney, liver) were subsequently dissected. Following isolation, all tissues were immediately snap-frozen on dry ice and then transferred to −80 °C for storage.

### Primary neuronal cultures

Primary motor neurons and hippocampal neurons were cultured as described previously [73, 83]. Taiwanese mice breeding pairs were set up as described above as timed matings to obtain heterozygous (*Smn*+*/-; SMN2tg/0*) and *Smn* null (*Smn-/-; SMN2tg/0*) embryos. At E13 (motor neurons) and E15 (hippocampal neurons), timed pregnant females were euthanized by cervical dislocation. Each embryo was processed individually and separately as genotyping was done retrospectively. For motor neurons, the ventral part of the spinal cord was dissected from each embryo, trypsinized using 0.05 % trypsin for 15 minutes at 37°C, treated with DNAsel (Sigma) and dissociated into a single cell suspension by trituration with a pipette. The cell suspension was separated using a 6.2% Optiprep gradient (Sigma) and centrifugation at 680 x *g* for 15 minutes. The motor neuron-containing fraction was pelleted and plated onto coverslips coated with poly-D,L-ornithine and laminin at a density of 10,000 cells per well in a 24 well plate. For mixed cultures of primary hippocampal neurons, the hippocampus was dissected and incubated in 0.25% trypsin for 15 min at 37°C after which the tissue was dissociated into single cells using fire-polished glass pipettes in culture medium (neurobasal medium supplemented with B27, glutamax, HEPES and penicillin/streptomycin). Hippocampal neurons were plated onto poly-D-lysine and laminin coated coverslips at a density of 20,000 cells per well in a 24-well plate. All primary cultures were grown in an incubator at 37°C and 5% CO_2_.

### Labeling of newly synthesized proteins in primary neurons

Novel protein synthesis in heterozygous control and SMA primary neurons was quantified using metabolic labeling with L-azidohomoalanine (AHA) using the Click-IT® AHA kit (ThermoFisher Scientific) according to the manufacturer’s recommendations. Briefly, after 7 days in culture, cells were washed with prewarmed PBS and the medium was replaced with methionine-free medium, supplemented with B27, Glutamax and l-azidohomoalanine and incubated for 30 minutes. The cells were subsequently fixed in prewarmed 4% PFA and permeabilized in 0.5% triton-X100 in PBS. A chemoselective ligation reaction between an azide and an alkyne allowed for subsequent labelling of newly synthesized proteins using Alexa Fluor 488. Cells were also labelled with nuclear marker DAPI. Primary neurons were imaged on a Zeiss LSM710 confocal microscope, where all healthy and morphologically normal neurons on a coverslip were imaged for subsequent analysis. The same confocal settings were used within each experiment for both SMA and control neurons. Images were subsequently analyzed in FIJI; the cell body was manually delineated and the mean fluorescence intensity was measured for each neuron. All analyses were performed with the operator blind to the genotype or treatment. The average fluorescence intensity was then calculated for heterozygous control and SMA motor neurons. As a control, the protein synthesis inhibitor anisomycin was used. Neurons were pre-treated with 40 µM anisomycin for 30 minutes, after which the medium was replaced with methionine-free medium as above, with the addition of 40 µM anisomycin. Cells were subsequently processed as described above.

### Polysome profiling

Cytoplasmic lysates from frozen mouse tissues were prepared as described previously [84]. Briefly, brains and spinal cords were pulverized in a pestle and mortar cooled by liquid nitrogen and the obtained powders were lysed in 0.8 mL of lysis buffer (10 mM Tris–HCl at pH 7.5, 10 mM NaCl, 10 mM MgCl2, 1% Triton-X100, 1% Na-deoxycholate, 600 U/ml RiboLock Rnase Inhibitor (Thermo Scientific), 1 mM DTT, 0.2 mg/ml cycloheximide, 5 U/ml Dnase I (Thermo Scientific)) by vigorous pipetting. Two consecutive centrifuges of 1 and 5 minutes respectively at 18,000 g at 4°C were performed to first remove cellular debris and then nuclei and mitochondria. The supernatants were immediately loaded on a linear sucrose gradient (15-50% sucrose [m/v], in 100 mM NaCl, 10 mM MgCl2, 10 mM Tris/HCl pH 7.5) and ultracentrifuged in a SW41Ti rotor (Beckman) for 1 h 40 min at 180,000 g at 4°C in a Beckman Optima™ LE-80K Ultracentrifuge. After ultracentrifugation, gradients were fractionated in 1 ml volume fractions with continuous monitoring absorbance at 254 nm using an ISCO UA-6 UV detector. Cytoplasmic lysates from cells were prepared as already described [40]. For EDTA treatment, the lysates were supplemented with 8 mM EDTA and left for 10 minutes on ice before being loaded on the gradients.

### Protein extraction from polysomal fractions and Western Blotting

Proteins were extracted from each sucrose fraction of the profile using the methanol/chloroform protocol [85]. Protein pellets were solubilized in Electrophoresis Sample Buffer (Santa Cruz Biotechnology) for the following SDS-PAGE and Western Blotting analysis. Proteins were separated in a 12% SDS-polyacrylamide gel or a 4-12% gradient gel (Novex) and blotted on nitrocellulose membrane or blotted using PVDF membrane stacks using the iBlot system (Life Technologies). Western blottings were performed using primary antibodies against RPL26 (1:2000, Abcam), RPS6 (1:1000, Cell Signalling), SMN (1:1000, BD Transduction Laboratories), hnRNP A1 (1:1000, Novus Biologicals), UBA6 (1:1000, Abcam) and the appropriate horseradish peroxidase (HRP) conjugated secondary antibodies (1:5000, Santa Cruz Biotechnology) or using the appropriate IRDye secondary antibodies (LICOR). Detection was performed using the ECL™ Prime Western Blotting Detection Reagent (Amersham) or when using infrared secondary antibodies, scanned on an Odyssey imager (LICOR) [86]. To quantify the co-sedimentation profiles of SMN along the sucrose gradient, we analysed Western Blots obtained using a semi-quantitative approach based on densitometric measurements (ImageJ). We calculated the percentage of the protein intensity of each fraction along the sucrose gradient as follows: %Pn = 100 * Dpn / sum Dpni, where Pn is the percentage of the protein in the fraction n; Dpn is the density of the protein in the fraction n and N is the total number of fractions. Western blots for NGS validation experiments were analysed using ImageStudio (LICOR).

### Analysis of the fraction of ribosomes on polysomes

The fraction of ribosomes on polysomes (FRP) was calculated from polysomal profiles as the ratio between the area under the curve of polysomes and the area under the curve of polysomes plus the area of the 80S peak. FRP was calculated for at least three different polysomal profiles (min 3 - max 9) for each tissue (spinal cord or brain), physiological condition (heterozygous control or SMA) and disease stage (pre-, early or late symptomatic). To bypass the problem of unclear discrimination of the 80S and polysomal peaks due to the sensitivity of absorbance along the sucrose gradient (for example, when the amount of tissue was limited), we first applied a blank profile subtraction. For each polysomal profiling run, 800 μL of lysis buffer was loaded on a gradient. The lysis buffer profile obtained was then subtracted from each tissue profile, which improved the final polysomal chromatogram and allowed for a more precise calculation of the FRP.

### Isolation of total and polysomal RNA and cDNA library preparation

RNA was extracted from control and SMA brains (at pre, early and late symptomatic stage), from ASO brains (at late symptomatic stage) and from control and SMA spinal cords (at late symptomatic stage). Profile fractions (total RNA) or polysomal fractions only (polysomal RNA) were pooled together. The fractions were first digested with proteinase K (100 mg/mL, Euroclone) for 90 minutes at 37°C and then RNA was extracted with acid-phenol:chloroform:isoamylalcohol, pH 4.5 (Sigma Aldrich) and isopropanol precipitation. cDNA libraries were produced starting from 500 ng of total or polysomal RNA (early and late symptomatic stage) using the TruSeq^®^ Stranded Total RNA (Illumina) kit according to manufacturer’s instructions.

### NGS data analysis

RNA-Seq and POL-Seq sequencing was performed with an Illumina HiSeq 2000 (*Mus musculus*, GPL13112). Fastq files were checked for quality control with FastQC. 100 bp reads generated from each sample were aligned to the mouse genome (GRCm38.p4) with Tophat (version 2.0.14), using the Gencode M6 transcript annotation as transcriptome guide. All programs were used with default settings unless otherwise specified. Mapped reads (ranging from 84% to 92% of total reads, Supplementary Table 1) were subsequently assembled into transcripts guided by reference annotation (Gencode M6) with Cufflinks (version 2.2.1). Expression levels were quantified by Cufflinks with normalized FPKM (fragments per kilobase of exon per million mapped fragments). Differentially expressed genes and transcripts in RNA-Seq and POL-Seq samples were detected with CuffDiff with a double threshold on the log2 fold change (absolute value > 0.75) and the corresponding statistical significance (p-value < 0.05). Translation efficiency analysis was performed with Xtail [54], genes showing differential translation efficiency values were selected using a threshold on Xtail p-value < 0.05. Functional annotation of gene lists and enrichment analysis with Gene Ontology terms and KEGG or REACTOME pathways were performed with the clusterProfiler Bioconductor package.

### qPCR

The retrotranscription reaction was performed starting from 1 μg of RNA using the RevertAid First Strand cDNA synthesis kit (Thermo Scientific) according to manufacturer’s protocol. qPCR was carried out using the CFX Connect™ Real-Time PCR Detection System (BioRad) using Kapa Syber Fast qPCR Mastermix (Kapa Biosystems) in 10 μL final reaction volume. Primer sequences are provided in Supplementary Table 2. Reaction conditions were as follows: one step of 95°C for 3 min and 40 cycles of 95°C for 2 sec of denaturation and 60°C for 25 sec of annealing and extension, followed by melting curve analysis (from 65 to 95 °C, increment of 0.5 °C every 0.5 sec). Actin and cyclophilin A were used as reference genes. Melting curves from each amplified product were analysed to ensure specific amplification. All reactions were performed in 2-4 biological replicates and 2-6 technical replicates. The obtained Ct values were used to calculate the fold change of each gene using the delta/delta Ct method [87]. The log2 delta TE (Translation Efficiency) was calculated as the difference between the fold change at the polysomal level and the fold change at the total level of the gene of interest.

### Electron microscopy

Tissue was prepared, processed and imaged using transmission electron microscopy as described previously [80].

### Statistical methods

For hierarchical clustering based on FRP, relative weight and relative righting time data were first rescaled in order for each variable to have the same mean and standard deviation. The distance matrix was calculated with the “euclidean” method. Clustering was performed with the “Ward.D” agglomeration method. Shifts in distribution mean values were tested with Student t-test. All other statistical analyses were performed in Graphpad Prism, with individual statistical tests used indicated in the figure legends.

## Acknowledgements

We thank the Core Facility, Next Generation Sequencing Facility (HTS) CIBIO for technical support, Prof. Paolo Macchi for the kind gift of the primary antibody against hnRNPA1; Efrem Bertini for technical support; Matteo Gaglio for skillful help with polysomal profiling and Hannah Shorrock for assistance with mouse breeding.

## Author contributions

P.B., F.M, EP and G.V. performed all polysomal purifications, RNA and protein extractions, western blotting and data analysis; T.T. performed all analyses of RNA-Seq and POL-Seq; E.G., H.N. and F.L. performed all mouse tissue collection, phenotypic data acquisition, primary neuron cultures and western blotting in Fig. 5d; P.Z. prepared the libraries of late-symptomatic samples; P.B. and E.P. prepared the libraries of early symptomatic samples; V.P. produced the SMN CRISPR/Cas9 cell-lines, H.Z. and F.M. performed ASO treatments in SMA mice; P.B. and E.P. performed qPCR experiments; P.B., T.T., E.P., E.G. prepare the figures; A.Q, T.G and G.V. conceived experiments and directed the research; P.B. T.T., E.G., F.L., A.Q, T.G and G.V wrote the manuscript. All authors contributed during preparation, revision and writing of the manuscript.

## Additional files

Additional File 1: Supplementary Methods, Supplementary Figures 1-9 and Supplementary Tables 1–2

Additional file 2: mice phenotypic and FRP data.

Additional file 3: translation efficiency data from RNA-Seq and POL-Seq performed on mice brains at early and late-symptomatic SMA stages.

Additional file 4: functional enrichment analysis on transcripts with altered translation efficiency values in SMA mice brains.

